# Designing of multiepitope-based vaccine against Leptospirosis using Immuno-Informatics approaches

**DOI:** 10.1101/2021.02.22.431920

**Authors:** Mohd Abdullah, Mohammad Kadivella, Rolee Sharma, Syed M. Faisal, Sarwar Azam

## Abstract

*Leptospira* is a zoonotic pathogen causing significant morbidity and mortality both in animals and humans. Although several surface proteins have been identified as vaccine candidate, they failed to induce sterilizing immunity and cross protection against different serovars. Thus, identification of highly immunogenic antigens that are conserved among pathogenic serovars would be first step towards development of universal vaccine for Leptospirosis. Here we used reverse vaccinology pipeline to screen core genome of pathogenic *Leptospira* spp.in order to identify suitable vaccine candidates. Based on properties like sub cellular localization, adhesin, homology to human proteins, antigenicity and allergenicity, 18 antigenic proteins were identified and were further investigated for immunological properties. Based on immunogenicity, Protegenicity, Antigenicity, B-cell and promiscuous T-cell epitopes, 6 Potential Vaccine Candidates (PVCs) were finally selected which covered most of the affected world population. For designing a Multi-Epitope Vaccine (MEV), 6 B-cell and 6 promiscuous MHC-I and MHC-II epitopes from each candidate were clustered with linkers in between and stitched along with a TLR4 adjuvant (APPHALS) at the N-terminal to form a construct of 361 amino acids. The physiochemical properties, secondary and tertiary structure analysis revealed that MEV was highly stable. Molecular docking analysis revealed the deep binding interactions of the MEV construct within the grooves of human TLR4 (4G8A). In-silico codon optimization and cloning of the vaccine construct assured good expression. Further, immune simulations have shown that MEV could induce strong and diverse B and T cell responses. Taken together our results indicate that the designed MEV could be a promising subunit vaccine candidate against Leptospirosis, however it requires experimental validation.

## 1.1 Introduction

Leptospirosis is a zoonosis caused by pathogenic *Leptospira* spp. It is one of the neglected and emerging tropical disease causing morbidity and mortality both in animals and humans (Sanchez-Montes 2015). In a recent report, average morbidity in human was estimated to be 1.03 million which causes nearly sixty thousand deaths across the world per year (Costa, Federico, et al. 2015, Sanchez-Montes 2015, Torgerson 2015). The disease has a huge impact on the health of companion and livestock animals causing serious economic consequences. The most viable strategy is vaccination to at-risk population of humans and host animals against leptospirosis. Currently available licensed vaccines for human and animals are in-activated whole cell vaccines which are also called bacterins. Few prominent leptospirosis vaccines in the market are SPIROLET from France (Bashiru 2018), Vax-Spiral from Cuba (Martínez, R. 2004, Bashiru 2018), bivalent vaccine from china (Hu 2014) and Nobivac L4 from UK (Bashiru 2018). However, These vaccines (bacterins) has limitation in its application due to numerous associated drawback such as lack of cross-reactivity or protection to only closely related serovars (Grassmann, 2017), lack of long term immunological memory, requirement of regular boosters due to short duration of immunity and sometimes high reactogenicity (Sonrier 2000, Dellagostin 2011, Dellagostin 2017, Adler 2015). Other factors which hamper the use of bacterins include high production costs and lack of consistency of production or repeatability (Dellagostin 2017, Adler 2015). Apart from these limitations there are high claims of death or transmission of infection in animals due lepto bacterins (Nobivac L4) (Klaasen 2013). These limitations and controversies emphasized on new approaches for leptospirosis vaccine design.

Emerging approaches to vaccine design and engineering are based on recent insights into immunology, structural biology and computational biology. These approaches are broadly termed as reverse vaccinology (RV) (Donati 2013). With the availability of whole genome sequences, reverse vaccinology can be applied more comprehensively and identify all potential antigen coded by a genome irrespective of their abundancy, phase of expression and immunogenicity (Pizza 2000, Seib 2012, Masignani 2002). Additionally, RV decreases the cost of vaccine production heavily and the method is more beneficial for pathogens like leptopspires which are slow growing in invitro (Guimarães, 2015). RV has been successfully applied to several pathogens, including serogroup B Neisseria meningitidis, Streptococcus agalactiae, Streptococcus pyogenes, Streptococcus pneumoniae and pathogenic Escherichia coli (Masignani 2019, Serruto, 2012). RV approaches select candidate proteins on basis of its structural features, such as: integral transmembrane, β-barrel arrangements, transmembrane alpha-helices, signal peptides, lipoproteins, secreted proteins, B-cell & T-cell epitopes. These candidate proteins are subjected to further development of recombinant protein vaccine or sub unit vaccine. Recently there has been a shift in RV approaches wherein multiple epitope based vaccines are designed rather than selecting the whole protein. Multi-eitope vaccine (MEV) are next generation vaccine design in which multiple epitopes from different proteins are stitched together using linkers and developed as subunit vaccine. For the last few years, this approach has shown its promising potential and many MEV vaccines are in clinical trials (Zhang, Lifang. 2018, Oyarzún et al. 2016). RV has been applied in *Leptospira* as well (Zeng, 2017, Grassmann, 2017, Dellagostin 2017). Most of the previous studies on *Leptospira* has been limited to identification of candidate proteins (like: Lp21,LP11, Lsa30, Lp35, OMPs,LIC20019,LIC20087 TonB-dependent receptor etc..,) and did not proceed towards development of multi-epitope vaccine (Felix 2020, Grassmann 2017). Many of the candidates (like: LP25, Lp21,LP11, Lsa30, Lp35,) which has been selected for validation and tested in animal model, failed to produce desired results (Felix 2020, Techawiwattanaboon 2019). However, RV has been evolving and getting more accurate with the availability of many new pathogenic genomes and development of latest bioinformatics and machine learning tools.

To overcome the short comings of previous research, in this study we reevaluated the genomes of all pathogenic species of *Leptospira* using RV approaches. Genes coding for proteins with adhesion like properties were screened and further assessed for B and T cell epitopes, promiscuous epitopes, population coverage and other Immunodominant properties such as antigenicity, immunogenicity and protegenicity (Scholzen et.al. 2019). Suitable vaccine candidates were selected on the basis of prominent immunological features. Finally, a Multiepitope vaccine (MEV) was designed using prospective epitopes from selected candidates and the construct was evaluated by downstream Insilco analysis for its feasibility/stability and immunodominant features.

## 2. Materials and Methods

### 2.1 Collection of proteins

In order to identify suitable vaccine candidates against *Leptopsira* through reverse vaccinology, proteome from soft-core genomes of pathogenic Leptospira was accessed from previous study (https://www.biorxiv.org/content/10.1101/2021.01.12.426470v2).

### 2.2 Prediction of adhesin like secretory or Outer-membrane proteins

Core proteome were screened for Outer-membrane (OM) and Extracellular (EC) protein using PSORTb v3.0.2 (https://www.psort.org/psortb/). The transmembrane topology was identified in each protein using TMHMM Server v. 2.0 (www.cbs.dtu.dk/services/TMHMM); proteins with multiple transmembrane spanning regions were ignored. Selected proteins were analyzed using SPAAN program (ref) for adhesins or adhesin like proteins.

### 2.3 Identification of host homologs

Adhesins similarity with host (i.e. human) proteins were checked using two methods (John 2012, Wilson 2000). First, adhesins were blasted with human protein dataset and were analyzed for micro level similarity with a cut-off of 35% identity in 80 amino acid length of the query (Silvanovich,2006). Second, blast between adhesins and host proteins should not have a continuous 9 or more identical matches in the alignment. Any protein that fails to qualify either of the criteria was discarded from the set of adhesins. Perl scripts were developed in-house to execute homology analysis and human protein dataset was downloaded from NCBI (https://www.ncbi.nlm.nih.gov/genome/guide/human/).

### 2.4 Antigenicity and allergenicity analysis

Selected adhesins were screened using an online version of VaxiJen v2.0 (Doytchinova 2007) to predict their antigenicity. The tool was run at a default parameter with a threshold value of 0.5. Highly antigenic adhesins were considered as potential protein antigens. Subsequently, the selected protein antigen which might cause allergenic action were identified by Allergen Online (http://www.allergenonline.org/databasefasta.shtml) database (Goodman 2016). Proteins which showed similarity in a sliding window of 80 amino acid segments with identities > 35% were labelled as potential allergen.

### 2.5 Protegenicity of protein antigen

Protective antigenicity of each protein antigen was predicted using Vaxigen-ML pipeline (http://www.violinet.org/vaxign/vaxign-ml/index.php). Vaxigen-ML provides score for each protein called as protegenicity score using an optimized supervised machine learning model. The tool uses supervised machine learning model which was based on training data set consisted of viral and bacterial antigens. We considered threshold protegenicity score of 90 to select effective antigens for vaccine development.

### 2.6 B-cell epitope prediction

B cell epitopes were predicted using BCPREDS software which contains two methods i.e. Amino Acid Pair (AAP) antigenicity method and subsequence kernel method (http://ailab-projects1.ist.psu.edu:8080/bcpred/predict.html). Epitopes (>=20 mers) with specificity >80% that resulted from each method were assembled and the consensus sequences were considered as the predicted B Cell epitopes (Sharma 2016). Further, these epitopes which are also predicted by Bepipred V2.0 from IEDB (>=20 amino acid thresholds) were selected as confident epitopes. Antigenicity of each confident epitope was calculated using Vaxijen v2.0 server. Epitopes with higher antigenicity were selected as final B cell epitopes.

### 2.7 T cell epitope, promiscuity and immunogenicity prediction

The epitopes of protein antigens binding to the reference set of MHC-I and MHC-II alleles were predicted using the Immune Epitope Database (IEDB) (Fleri et al., 2017) online server (MHC-I version: 2.23 and MHC-II version: 2.22), using the IEDB-recommended parameters. More specifically, stringent consensus percentile ranking scores cut-off of < 0.1% was used to select MHC-I epitopes, while consensus percentile ranking scores cut off of < 1% was used for MHC-II strong binders. Additionally, epitopes which restricts more than 4 allele of HLA-I or HLA-II in the reference set (Davila et al., 2012; McNamara et al., 2010) were screened as Promiscuous epitopes using perl script. Immunogenicity score was calculated for each MHC-I epitope reflecting the immunogenicity of the peptide-MHC (pMHC) complex formation, using IEDB tool “T cell class I pMHC immunogenicity predictor” on default parameter (Calis et al., 2013). Immunogenicity of each protein was determined by summing the immunogenicity scores of all the 9-mer epitopes of each protein predicted to bind the MHC-I reference set of alleles (ong et al 2020).

### 2.8 Estimation of population coverage

The population coverage of each protein antigen were determined using standalone IEDB Population Coverage Tool v1.0.1(Bui et al., 2006). MHC-I and MHC-II reference alleles interacting to the epitopes of each protein antigen were together subjected as input to IEDB population coverage tool. We calculated population coverage for the whole world as well as south Asian countries like India, Pakistan, Sri Lanka and Bangladesh as these are having high disease burden as per WHO report.

### 2.9 Immune-simulation study of selected vaccine candidate genes

Immune response profiles of vaccine candidate genes were characterized by C-ImmSim server (http://150.146.2.1/C-IMMSIM/index.php) under default parameter (Rapin et al., 2010). Server uses machine learning methods and position-specific scoring matrices to identify the epitope peptides and other immune interactions.

### 2.10 Novel Multi Epitope Vaccine Construct designing and evaluation

Promiscuous MHC-I, MHC-II and B cell epitopes of 6 potential vaccine candidates were clustered using “Epitope Cluster Analysis Tool” (http://tools.iedb.org/cluster/) (Ullah et.al. 2020) Epitopes were clustered on sequence identity threshold of 70%. Epitopes were selected for multi epitope vaccine design in a way that it represented at least one epitope for MHC-I, MHC-II and B cell from each vaccine candidates. B-cell epitope, MHC-I epitopes and MHC-II epitopes were selected on antigenicity, immunogenicity and promiscuity respectively. Overall, overlapped epitopes were preferred in each class (Rahman et al 2020). Hydropathy Index of each epitopes were calculated using GRAVY analysis from ProtParam (https://web.expasy.org/protparam/). Epitopes were sorted on hydropathy index and linked together using three amino linkers that is “GGGGS”, “GGGS”, “GPGPG” for B cell, MHC-I and MHC-II epitopes respectively. Subsequently, EAAAK was used to join “APPHALS”, a TLR-4 adjuvant at the N-terminal of Multi-epitope vaccine construct (Chaudhuri et al. 2020). MEV was evaluated for different subunit vaccine properties. Accordingly, different physiochemical properties such as molecular weight (MW), theoretical isoelectric point (pI), half-life, aliphatic index, GRAVY were calculated using ProtParam and solubility was assessed by SOLpro. Allergenicity and Antiinflammatory properties were evaluated using AllergenOnline and AIPpred respectively. Antigenicity was estimated by Vaxijen v2.0 and Immune response profiles were generated by C-ImmSim server (Rapin et.al 2006).

### 2.11 Secondary and tertiary structure prediction of MEV and its refinement

Secondary structure including alpha helix, beta sheet and turns region of MEV was characterized by Chou and Fasman Secondary Structure Prediction (CFSSP) server (Kumar et.al 2013) (http://www.biogem.org/tool/chou-fasman/index.php) and solvent accessibility of each amino acid residue was assessed using RaptorX server (http://raptorx.uchicago.edu/). RaptorX server was used to predict Novel MEV’s 3D structure (Källberg et.al. 2014). Predicted 3D structure were further refined by ModRefiner (https://zhanglab.ccmb.med.umich.edu/ModRefiner/) to improve the accuracy. Refined model structure was validated by Ramachandran plot using Procheck server (SAVES v6.0, https://saves.mbi.ucla.edu/) and ProSA-web (https://prosa.services.came.sbg.ac.at/prosa.php) server.

### 2.12 Docking with TLR & MD simulation of docked complex

The 3D structure of novel MEV protein was docked with TLR4 (PDB ID: 4G8A) receptors using Patchdock web server (https://bioinfo3d.cs.tau.ac.il/PatchDock/) (Schneidman-Duhovny et.al 2005). The Docked complex was further refined using FireDock (http://bioinfo3d.cs.tau.ac.il/FireDock/) (Mashiach et.al 2008). Intramolecular interactions of the protein-protein docked complex were visualized using DIMPLOT program of the LigPlot+ and UCSF Chimera version 1.15 (Pettersen et.al 2004). Molecular dynamics simulations were performed using GROMACS 2020.5 package to evaluate the stability, energy minimization of TLR4-MD2-MEV complex, having a lowest binding energy (Abraham et.al 2015). OPLS-AA force field (Kaminski et.al 2001) and TIP3P water model (Mark et.al 2001) were used in the simulation. Further, a cubic box of 20Å distance from any edge of the box along with spc216 were used to solvate the system. In the next step, the system was neutralized by adding 21 sodium ions. Energy minimization of the neutralized system was executed by employing the steepest descent algorithm until the maximum force was less than 1000.0 kJ/mol/nm. After the minimization process, NVT equilibration for 100ps at 300K and NPT ensemble equilibration for 100ps at 1 bar pressure were performed for each system. The final MD production run were performed for 1ns with the maximum number of steps of 500000. In order to analyse the output files VMD, Xmgrace and other gromacs inbuilt tools were used.

### 2.14 Codon optimization and In-silico cloning of MEV

Codon of MEV was optimized for *E. coli* K12 strain using JCat (http://www.prodoric.de/JCat) server. The server was run with criteria (i) avoid termination of rho-independent transcription, (ii) Avoid binding-site of prokaryote ribosome, and (iii) Avoid cleavage-sites of restriction enzymes for BamHI and XhoI. SnapGene software (https://www.snapgene.com/) was used to insert optimized codon sequence into the E. coli pET28a (+) vector between restriction sites of XhoI and Bam HI at multiple cloning site. In silico expression of the peptide was visualized using SnapGene software.

## Results

### Proteome dataset in the study

In previous study, Comparative genomics of all the sequenced Leptospira genomes established that 26 species are highly pathogenic in nature (https://www.biorxiv.org/content/10.1101/2021.01.12.426470v2). Analysis of pathogenic Leptospira revealed that soft-core genome consists of 2408 genes of which 1478 genes are present in all (100%) species whereas rest of the genes are present in at least 95% of species. Softcore genome i.e. 2408 genes are highly conserved and they were taken as initial dataset for RV workflow (Figure 1) in this study.

**Figure 1:**
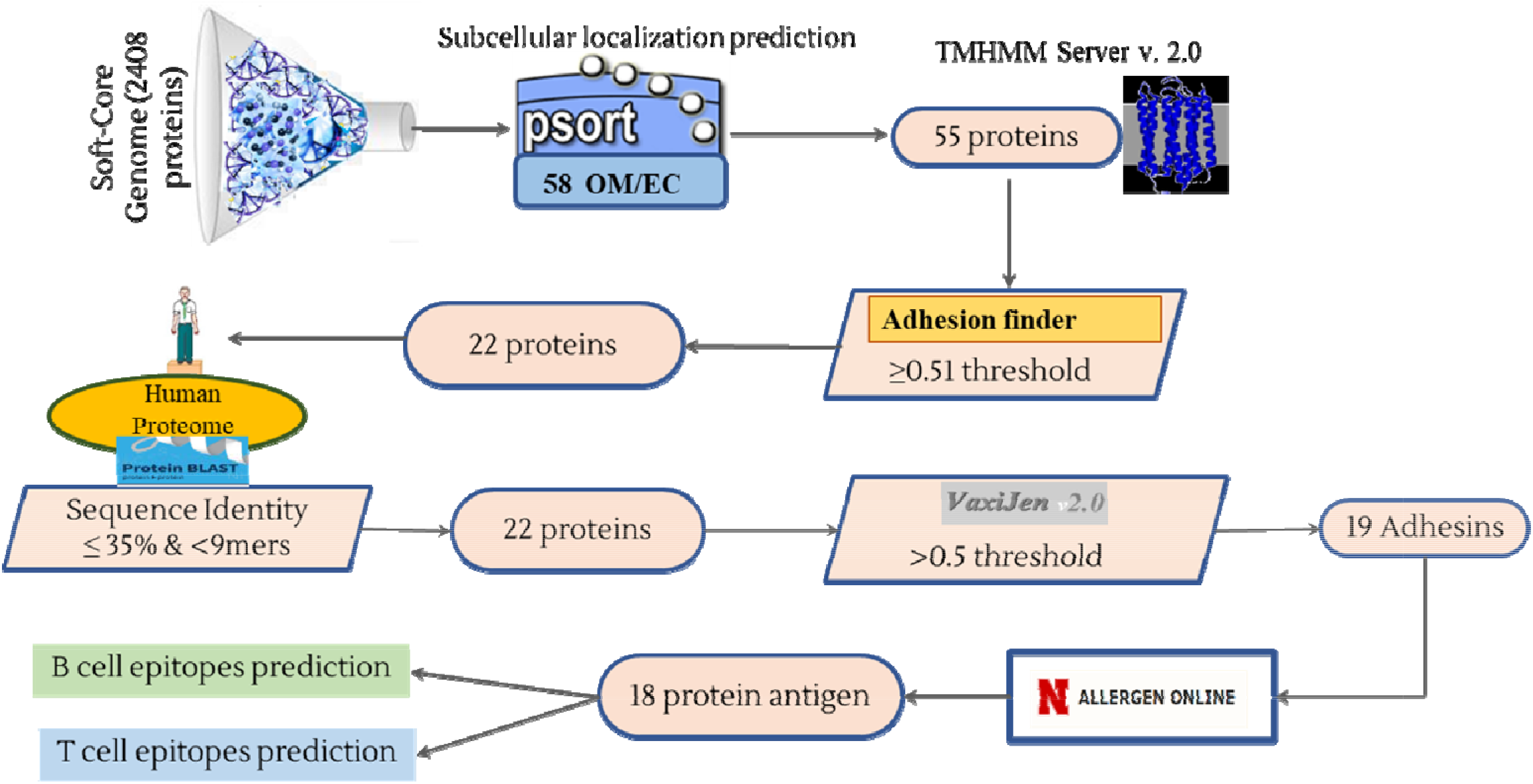
Schematic representation of Reverse vaccinology pipeline

### Prediction of adhesin like proteins

Soft-core genome analysis for subcellular localization predicted 45 proteins to be outer membrane and 13 proteins to be secretory in pathogenic Leptospira (Figure 2) & (Supplementary Table 1). Collectively, these 58 proteins were further screened for transmembrane helices. Three proteins having more than one trans membrane helices, were discarded from dataset. Selected 55 proteins with one or zero transmembrane helices were analyzed for adhesin capability (Supplementary Table 2). Highest adhesin probability observed was 0.956 whereas lowest was 0.157. Proteins with credible adhesin like properties were filtered on a cutoff of 0.51 and 22 proteins were considered as adhesins. None of the selected adhesins were having homology or identity with human proteome. Signal sequences of each proteins were removed and only functional sequence were regarded as potential antigen and subjected for downstream analysis. Summary of features of each adhesins are provided in Table 1.

**Figure 2:**
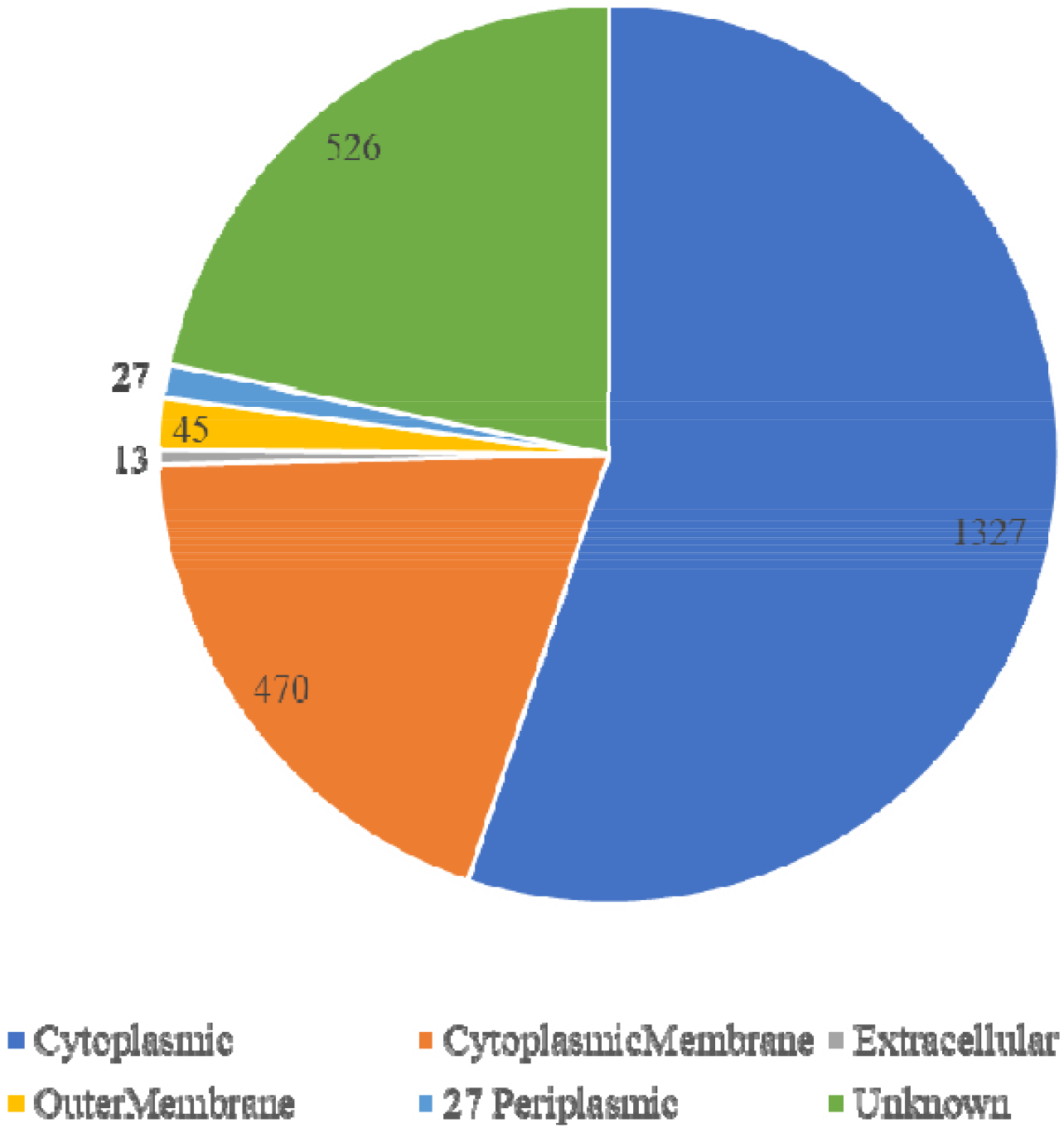
Sub cellular localization of soft core genome of pathogenic *Leptospira*

**Table 1:** Summary of features of 19 Adhesins identified in the study.

| S. No. | Protein Accession | Protein Length | Localization | Localization Probability | Adhesin Probability | Trans-membrane helices | Antigenicity |
| --- | --- | --- | --- | --- | --- | --- | --- |
| 1 | WP_011670651.1 | 235 | Extracellular | 0.965 | 0.921 | 0 | 0.6718 |
| 2 | WP_020780498.1 | 234 | Extracellular | 0.965 | 0.638 | 0 | 0.8016 |
| 3 | NP_714239.2 | 422 | Outer Membrane | 0.949 | 0.631 | 1 | 0.7549 |
| 4 | WP_011671354.1 | 230 | Extracellular | 0.965 | 0.941 | 0 | 0.6394 |
| 5 | WP_011669418.1 | 209 | Extracellular | 0.964 | 0.749 | 1 | 0.9531 |
| 6 | WP_011669449.1 | 252 | Extracellular | 0.964 | 0.956 | 0 | 0.9421 |
| 7 | WP_011670794.1 | 234 | Outer Membrane | 0.949 | 0.548 | 1 | 0.8778 |
| 8 | NP_712625.2 | 323 | Outer Membrane | 0.949 | 0.688 | 0 | 0.6047 |
| 9 | **WP_011671323.1** | **800** | **Outer Membrane** | **0.952** | **0.547** | **0** | **0.5494** |
| 10 | WP_011671327.1 | 867 | Outer Membrane | 0.949 | 0.693 | 1 | 0.6058 |
| 11 | WP_011669213.1 | 647 | Outer Membrane | 0.949 | 0.764 | 1 | 0.6418 |
| 12 | WP_011669397.1 | 426 | Outer Membrane | 0.949 | 0.636 | 0 | 0.6195 |
| 13 | WP_011669637.1 | 438 | Outer Membrane | 1 | 0.822 | 0 | 0.6665 |
| 14 | WP_011670051.1 | 547 | Outer Membrane | 0.949 | 0.651 | 1 | 0.5982 |
| 15 | WP_011670465.1 | 610 | Outer Membrane | 0.949 | 0.808 | 0 | 0.7256 |
| 16 | WP_011670696.1 | 464 | Extracellular | 1 | 0.729 | 0 | 0.7212 |
| 17 | WP_011670788.1 | 486 | Outer Membrane | 0.949 | 0.774 | 0 | 0.6474 |
| 18 | WP_011670856.1 | 777 | Outer Membrane | 1 | 0.544 | 1 | 0.5831 |
| 19 | WP_011670925.1 | 472 | Extracellular | 0.964 | 0.843 | 0 | 0.5072 |

### Antigenicity, protegenicity and allergenicity analysis

Adhesins with higher antigenicity are considered as better protein antigen for vaccine candidate (Monterrubio-López et.al 2015, Rashid et.al 2019). While screening for antigenicity, 19 adhesins out of 22 proteins with VaxiJen score more than 0.5 were considered as prospective protein antigens (Supplementary Table 3). Further, protegenicity (protective antigenicity) for these 19 adhesins were evaluated using recently developed Machine learning based pipeline by Ong et al. (Ong et al. 2020). A protegenicity score cutt-off of 90 can be safely considered as criteria to mine out good antigens for vaccine development. In our analysis, all adhesins were predicted with protegenicity score more than 90 (Table 2). Thus, each protein can be considered as a suitable antigen for vaccine candidate on the basis of protegenicity score. These 19 proteins were then screened for allergenicity in human. Only one Outer Membrane protein (WP_011671323) showed significant match with known allergen “Sar s 1 allergen Yv5032C08” and it was excluded (Table 1). Finally, the resultant 18 protein antigens were analyzed for topnotch potential vaccine candidates (PVC) in accordance with their immunological properties.

**Table 2:** Adhesin like proteins has been predicted to induce protective Immunity.

| **S. No** | **Protein Accession ID** | **Protegenicity (%)** |
| --- | --- | --- |
| 1 | WP_011671354.1 | 90.909091 |
| 2 | WP_011669449.1 | 90.909091 |
| 3 | WP_011670925.1 | 90.909091 |
| 4 | WP_011670651.1 | 91.000687 |
| 5 | NP_712625.2 | 92.305931 |
| 6 | WP_020780498.1 | 92.534921 |
| 7 | WP_011670794.1 | 93.771468 |
| 8 | WP_011669418.1 | 93.863064 |
| 9 | NP_714239.2 | 93.977559 |
| 10 | WP_011670788.1 | 94.023357 |
| 11 | WP_011669397.1 | 94.916419 |
| 12 | WP_011670051.1 | 97.984887 |
| 13 | WP_011669637.1 | 98.030685 |
| 14 | WP_011670856.1 | 98.419968 |
| 15 | WP_011670696.1 | 99.404626 |
| 16 | WP_011670465.1 | 99.679414 |
| 17 | WP_011669213.1 | 99.725212 |
| 18 | WP_011671327.1 | 99.748111 |

### B cell epitope Identification

B cell epitope in all the potential protein antigens were identified using two different tools i.e. BCPREDs and Bepipred. A total of 181 epitopes were identified using BCPREDS tool in all the 18 proteins antigen with length ranging from 20 to 65 amino acids (Supplementary Table 4). Further, 32 epitopes having low antigenicity were discarded and the remaining 149 were taken as BCPRED’s predicted epitopes. Another tool, IEDB Bepipred provides score for each amino acids of a protein. Bepipred output of each protein has been summarized in Figure 3. Amino acids with scores above threshold value are depicted as yellow in the figure and can be regarded as epitopic in nature. BCPred’s Predicted epitopes which were also described by Bepipred either identical or with sufficient overlap, were selected as confident B cell epitopes. A confident set of 72 epitopes from all the 18 protein antigen has been provided in (Table 3). Each protein consists of at least one confident B-cell epitope whereas protein “WP_011671327” and “WP_011670696” have maximum of 7 epitopes (Supplementary Table 5).

**Figure 3.**
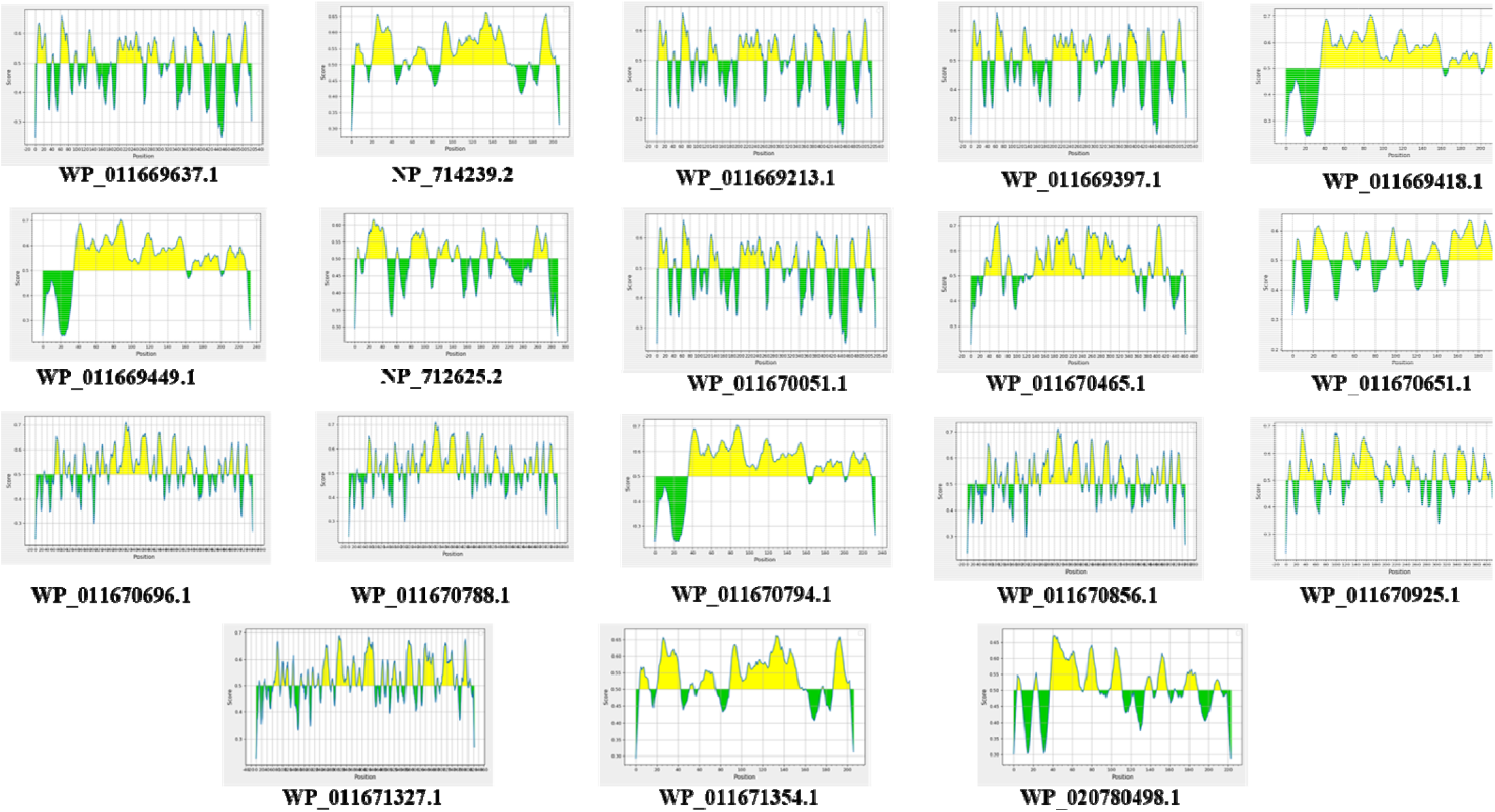
Bepipred output of each Protein Antigen: Amino acid with score above threshold value are depicted as in yellow in the figure.

**Table 3:**
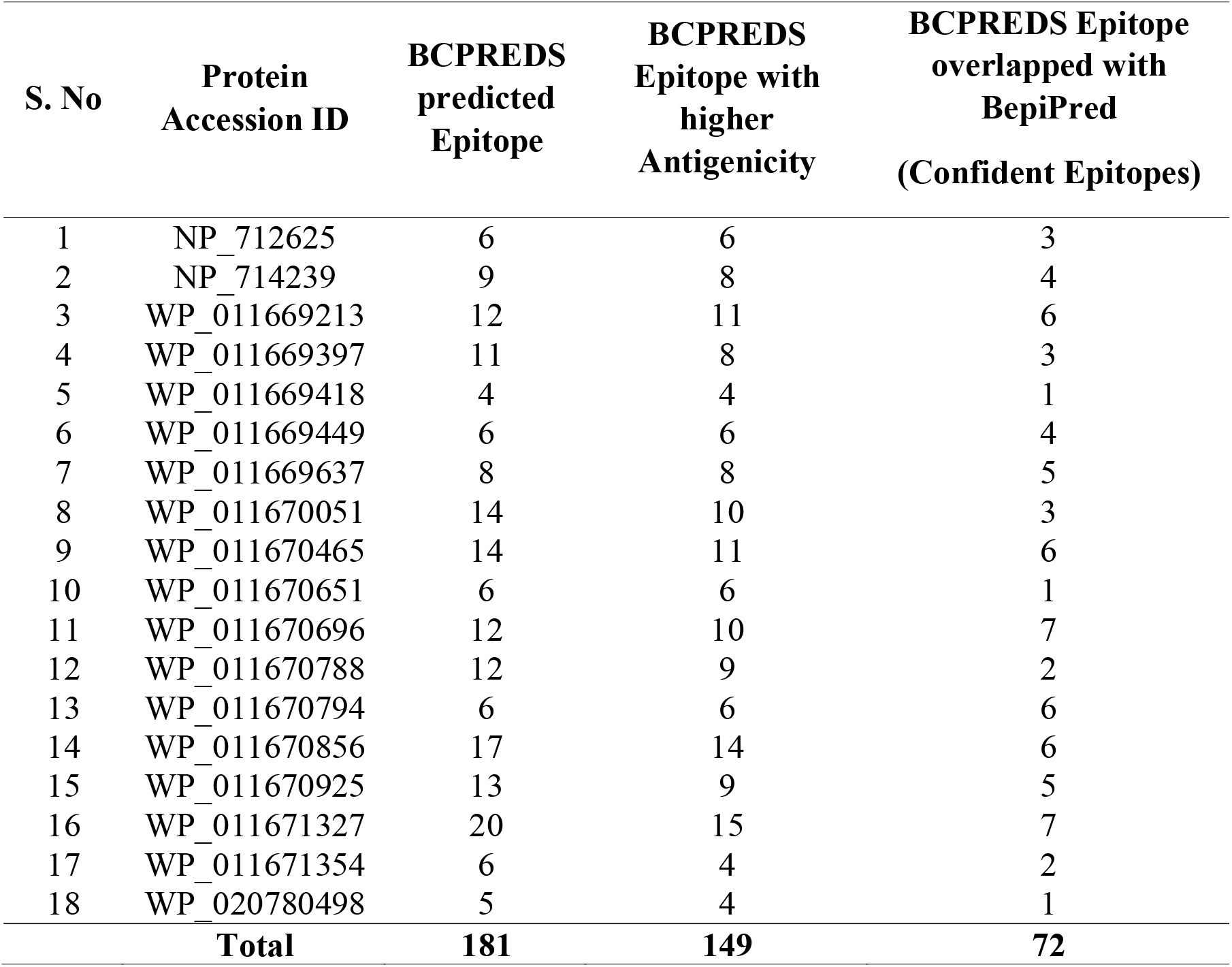
Prediction of B cell Epitopes by two tools BCPREDS and Bepipred.

**Immunity.**
| **S. No** | **Protein**<br>**Accession ID** | **BCPREDS**<br>**predicted**<br>**Epitope** | **BCPREDS**<br>**Epitope with**<br>**higher**<br>**Antigenicity** | **BCPREDS Epitope**<br>**overlapped with**<br>**BepiPred**<br>**(Confident Epitopes)** |
| --- | --- | --- | --- | --- |
| 1 | NP_712625 | 6 | 6 | 3 |
| 2 | NP_714239 | 9 | 8 | 4 |
| 3 | WP_011669213 | 12 | 11 | 6 |
| 4 | WP_011669397 | 11 | 8 | 3 |
| 5 | WP_011669418 | 4 | 4 | 1 |
| 6 | WP_011669449 | 6 | 6 | 4 |
| 7 | WP_011669637 | 8 | 8 | 5 |
| 8 | WP_011670051 | 14 | 10 | 3 |
| 9 | WP_011670465 | 14 | 11 | 6 |

**Table 3 : Prediction of B cell Epitopes by two tools BCPREDS and Bepipred.**
|  |  |  |  |  |
| --- | --- | --- | --- | --- |
| 10 | WP_011670651 | 6 | 6 | 1 |
| 11 | WP_011670696 | 12 | 10 | 7 |
| 12 | WP_011670788 | 12 | 9 | 2 |
| 13 | WP_011670794 | 6 | 6 | 6 |
| 14 | WP_011670856 | 17 | 14 | 6 |
| 15 | WP_011670925 | 13 | 9 | 5 |
| 16 | WP_011671327 | 20 | 15 | 7 |
| 17 | WP_011671354 | 6 | 4 | 2 |
| 18 | WP_020780498 | 5 | 4 | 1 |
| **Total** |  | **181** | **149** | **72** |

### Promiscuity and immunogenicity of T cell epitope

T cell epitopes which binds strongly to MHC allele can cause MHC restriction. We identified such epitopes binding to MHC-I reference allele in the range of 10 to 94 for each of 18 antigens. Similarly, epitopes binding to MHC-II reference alleles were predicted in the range of 15 to 87 for each antigen. Out of 18 antigens investigated, only two were having epitopes which can bind to all 27 reference alleles of MHC-I (Table 4). On the other hand, two antigen (“NP714239”, “wp0116713 27”) were having epitopes which can bind to maximum of 19 reference alleles of MHC-II. Taken together, antigen “wp011671327” was predicted to have largest number of T cell epitopes (168) and binds to all MHC-I reference allele and 19 MHC-II reference alleles. The predicted epitopes were filtered for promiscuous epitopes on the basis of its binding to four or more MHC-I and MHC-II reference set of alleles. A total of 69 epitopes were identified as promiscuous epitopes of which 29 epitopes binds to MHC-I reference allele and 40 epitopes binds to MHC-II reference alleles (Supplementary Table 6, Supplementary Table 7). Out of total 18 protein antigens, 13 proteins were predicted to have promiscuous epitopes for MHC-I reference alleles whereas only 8 proteins were predicted to have promiscuous epitopes for MHC-II reference alleles. However, only seven protein antigens were predicted to have promiscuous epitopes for both MHC-I and MHC-II reference set of alleles. Maximum number of MHC-I promiscuous epitopes (4) were found in antigen “NP_712625” which binds to 13 alleles of the reference set (Table 4 and Supplementary Table 6) whereas antigen “WP_011670465” consists a maximum of 10 promiscuous epitopes of MHC-II (Table 4 & Supplementary Table 7).

**Table 4:** Immunogenicity and the total number of predicted epitopes and binding alleles for MHC-I and MHC-II reference alleles of 18 potential antigens.

| S. No. | Protein Accession ID | Immunogenicity Score | MHC-I Reference alleles |  | MHC-II Reference alleles |  |
| --- | --- | --- | --- | --- | --- | --- |
|  |  |  | Total Predicted Epitopes | Predicted Binding Alleles/Total alleles | Total Predicted Epitopes | Predicted Binding Alleles/Total alleles |
| 1 | NP_712625 | 1.51444 | 48 | 23/27 | 69 | 13/27 |
| 2 | NP_714239 | 1.63616 | 53 | 24/27 | 79 | 19/27 |
| 3 | WP_011669213 | 0.27117 | 62 | 25/27 | 87 | 16/27 |
| 4 | WP_011669397 | 1.01257 | 47 | 24/27 | 55 | 7/27 |
| 5 | WP_011669418 | -0.22037 | 11 | 10/27 | 15 | 7/27 |
| 6 | WP_011669449 | -0.13648 | 13 | 12/27 | 25 | 4/27 |
| 7 | WP_011669637 | 0.65048 | 26 | 22/27 | 40 | 13/27 |
| 8 | WP_011670051 | 2.53251 | 61 | 27/27 | 69 | 15/27 |
| 9 | WP_011670465 | 0.58537 | 52 | 25/27 | 59 | 15/27 |
| 10 | WP_011670651 | -0.16725 | 10 | 16/27 | 26 | 5/27 |
| 11 | WP_011670696 | 0.61363 | 26 | 22/27 | 50 | 5/27 |
| 12 | WP_011670788 | 1.59892 | 32 | 23/27 | 30 | 8/27 |
| 13 | WP_011670794 | -0.81439 | 14 | 13/27 | 38 | 12/27 |
| 14 | WP_011670856 | -0.94868 | 58 | 26/27 | 70 | 14/27 |
| 15 | WP_011670925 | 1.9916 | 36 | 20/27 | 35 | 8/27 |
| 16 | WP_011671327 | 4.4854 | 94 | 27/27 | 74 | 19/27 |
| 17 | WP_011671354 | 0.59313 | 21 | 15/27 | 42 | 8/27 |
| 18 | WP_020780498 | -1.65195 | 18 | 21/27 | 21 | 5/27 |

Calculation of immunogenicity score for 18 antigens showed that only 12 antigens are immunogenic with positive score (Supplementary Table 8). Among 12 antigens, it is observed that antigen “wp011671327” was most immunogenic followed by antigen WP_011670051, WP_011670925, NP_714239, WP_011670788, NP_712625, WP_011669397, WP_011669637, WP_011670696, WP_011671354, WP_011670465 and WP_011669213. Six out of seven proteins which have promiscuous epitopes for both MHC-I and MHC-II reference set of alleles are also noticed to be highly immunogenic with positive score.

### Population Coverage Analysis

Population coverage is the assessment of individuals in the population which can interact with epitopes of antigen. Antigen “NP_714239” and “WP_011671327” was predicted with 100 % population coverage over the World and South Asian population and thus having highest coverage among all 18 antigens (Table 5 & Figure 4). Population Coverage of remaining 16 antigens was obtained in the range of 99.99% to 92.31% for World and 99.99% to 91.04% for South Asian countries. The antigen “WP_011669449” had lowest predicted coverage for each population i.e. World and South Asia.

**Figure 4.**
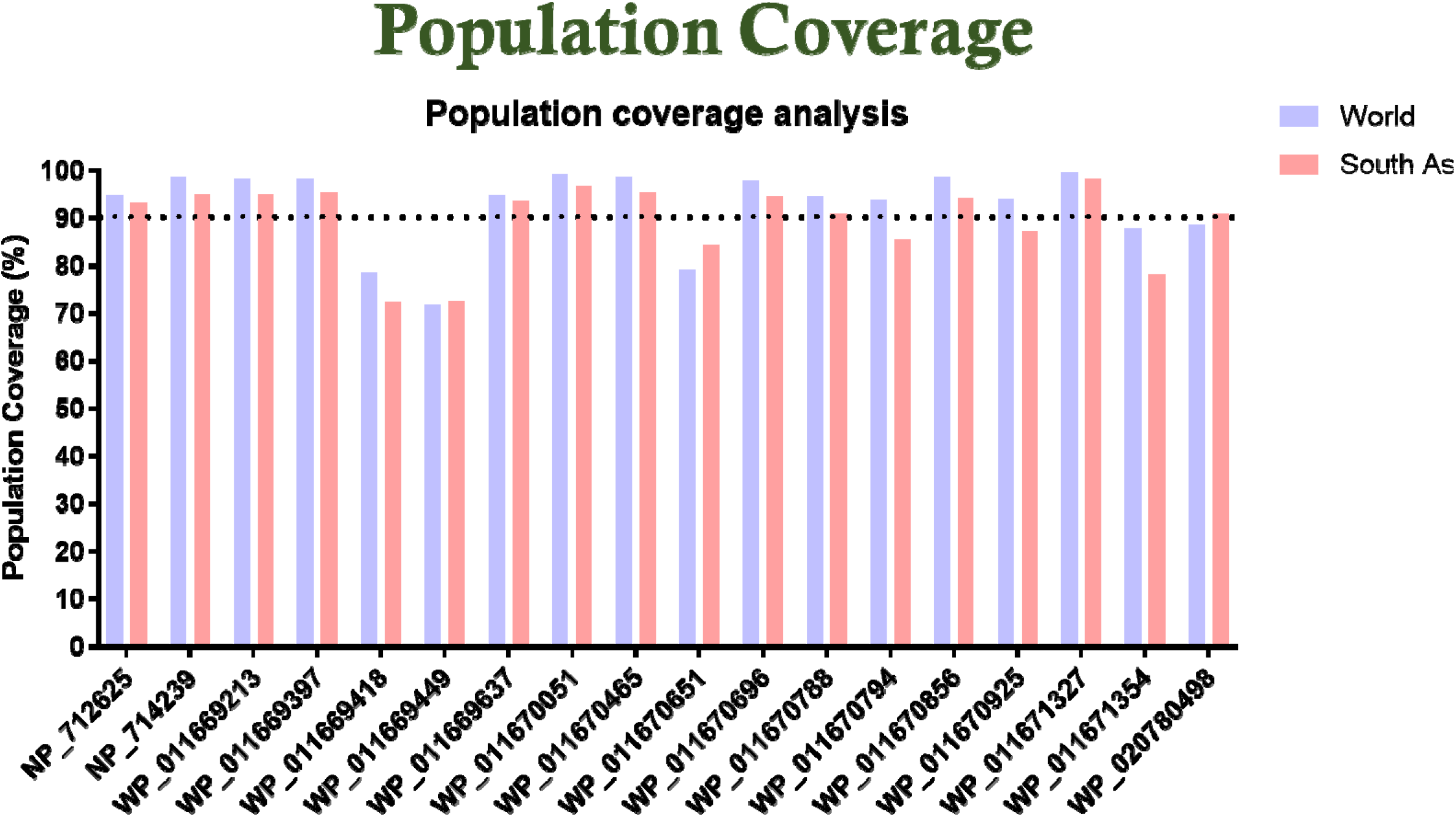
Population coverage of 18 antigens by IEBD standalone population coverage.

**Table 5:**
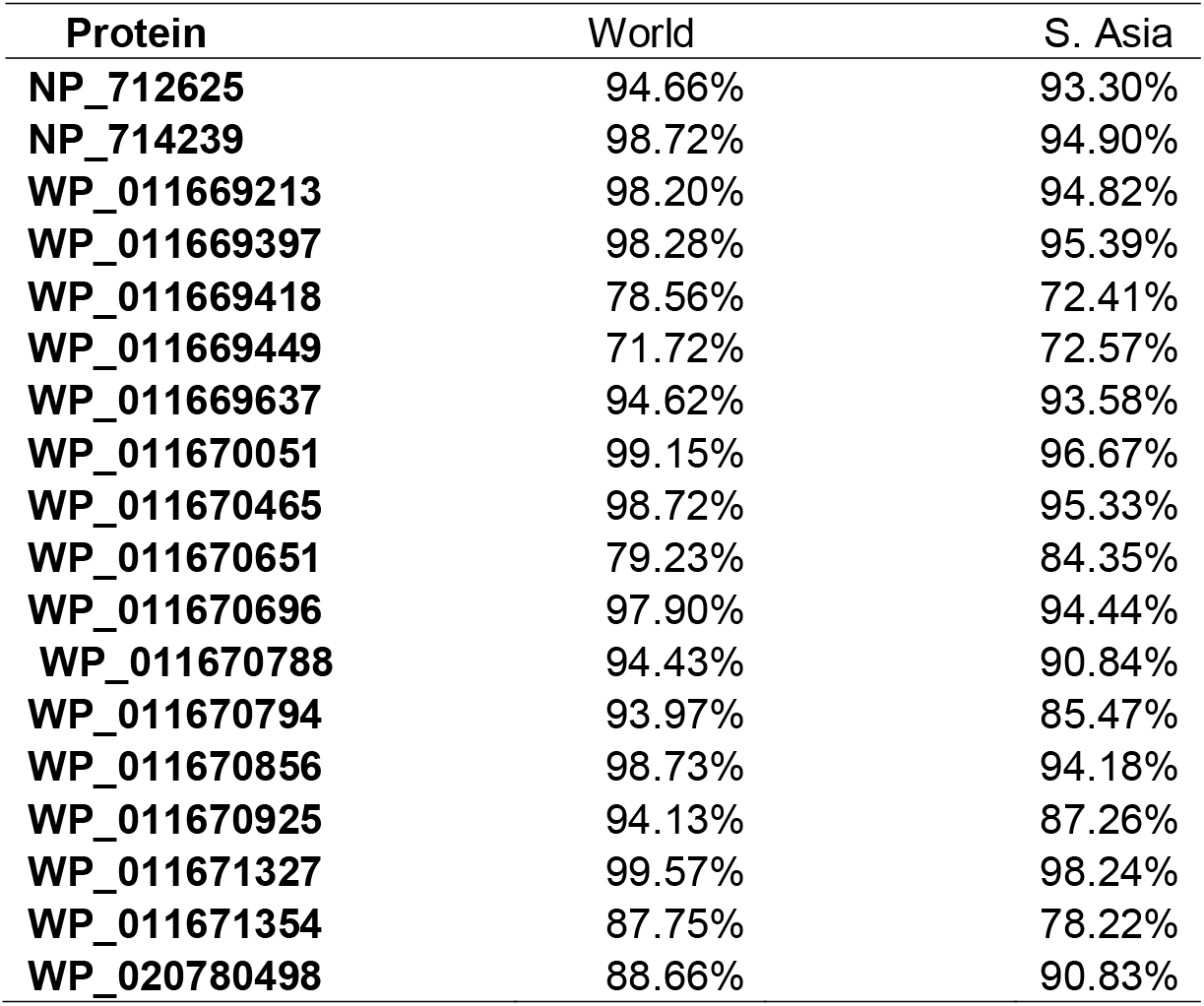
Population coverage analysis of 18 protein antigens over the World and South Asian population.

| **Protein** | **World** | **S. Asia** |
| --- | --- | --- |
| **NP_712625** | 94.66% | 93.30% |
| **NP_714239** | 98.72% | 94.90% |
| **WP_011669213** | 98.20% | 94.82% |
| **WP_011669397** | 98.28% | 95.39% |
| **WP_011669418** | 78.56% | 72.41% |
| **WP_011669449** | 71.72% | 72.57% |
| **WP_011669637** | 94.62% | 93.58% |
| **WP_011670051** | 99.15% | 96.67% |
| **WP_011670465** | 98.72% | 95.33% |
| **WP_011670651** | 79.23% | 84.35% |
| **WP_011670696** | 97.90% | 94.44% |
| **WP_011670788** | 94.43% | 90.84% |
| **WP_011670794** | 93.97% | 85.47% |
| **WP_011670856** | 98.73% | 94.18% |
| **WP_011670925** | 94.13% | 87.26% |
| **WP_011671327** | 99.57% | 98.24% |
| **WP_011671354** | 87.75% | 78.22% |
| **WP_020780498** | 88.66% | 90.83% |

### Selection of Potentials Vaccine Candidates (PVC)

Reverse vaccinology pipeline was applied to select adhesins as potential vaccine candidates. Initially strong 18 protein antigens were identified on the basis of antigenicity, protegenicity and allergenicity. These antigens were then strictly assessed on different immunogenic parameters for their selection as potential vaccine candidate. Abundance of B cell and T cell epitopes, availability of promiscuous epitopes, immunogenicity and population coverage of each antigen were evaluated. A non-redundant set of 6 antigens which qualify all the parameters were selected as final vaccine candidates (Table 6). These PVCs can be used to develop monovalent or multivalent vaccine for leptospirosis.

**Table 6:** Final 6 Potential Vaccine Candidates that qualified all the parameters against *Leptospirosis.*

| S. No | Accession ID | Immunogenicity<br>MHC-I | No. of promiscuous<br>Epitopes |  | B-Cell |
| --- | --- | --- | --- | --- | --- |
|  |  |  | MHC-I | MHC-II |  |
| 1 | NP_712625 | 1.51444 | 4 | 1 | 4 |
| 2 | NP_714239 | 1.63616 | 3 | 7 | 4 |
| 3 | WP_011669637 | 0.65048 | 1 | 4 | 5 |
| 4 | WP_011670051 | 2.53251 | 2 | 5 | 3 |
| 5 | WP_011670465 | 0.58537 | 3 | 10 | 6 |
| 6 | WP_011671327 | 4.4854 | 1 | 8 | 7 |

### Novel Multi Epitope Vaccine Construct designing and evaluation

We further improvise our study to design a Multi Epitope Vaccine (MEV) to ensure better immunity by combining prominent epitopes of all the 6 PVCs. Initially, a total of 77 epitopes from all the six PVCs which contain all B cell epitopes and promiscuous epitopes of T-cell were clustered into 48 epitopes including 39 singletons (Supplementary Table 9). As described in the method, a total of 15 epitopes were strategically selected and stitched together with linkers into one MEV construct. The length of MEV construct was 361 amino acids which includes APPHALS adjuvant as well (Figure 5 (A) & (B)). The MEV construct has a molecular weight of 37.8 kDa, a theoretical PI of 4.92 and an instability index of 31.14. The construct is thermostable and non-polar with an aliphatic coefficient of 55.46 and GRAVY value of −0.384. The multi epitope protein will be soluble upon over expression of the construct as the predicted solubility score is 0.694. The construct was found to be non-allergic, anti-inflammatory and highly antigenic in nature with an antigenicity score of 1.3082. The aforementioned analysis suggests that the MEV is a highly suitable subunit vaccine candidate (Supplementary Table 10).

**Figure 5:**
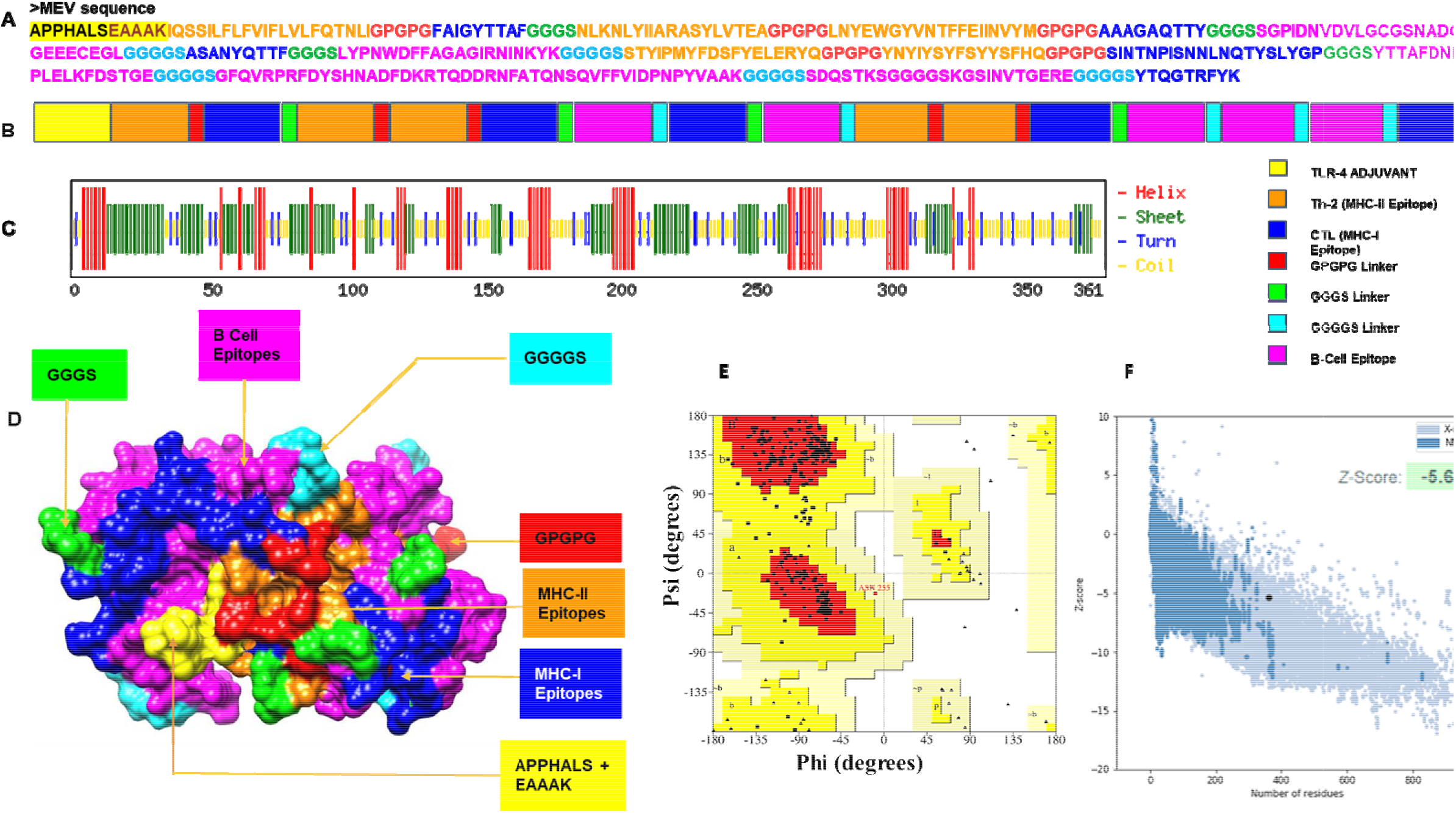
Design, construction and structural validation of multi-epitope vaccine candidate for Leptospirosis. (A) The predicted primary structure of the MEV with adjuvant APPHALS and linkers GPGPG, GGGS and GGGGS together with respective T cell and B cell epitopes. (B) Structural domains and epitopes rearrangement of MEV according to the sequence in A. (C) Secondary structure of MEV as analyzed through CFSSP: Chou and Fasman secondary structure prediction server (D) Final refined tertiary structure of MEV (surface view) visualised in UCSF Chimera; domains and epitopes are represented in different colors (APPHALS+EAAAK: Yellow, B cell epitopes: Magenta, MHC I Epitopes: Blue, MHC II Epitopes: Orange, Linkers: GPGPG: Red, GGGS: Green, GGGGS: Cyan). (E) validation of the refined model with Ramachandran plot analysis showing 90.4%, 9.3% and 0% of protein residues in favored, allowed, and disallowed (outlier) regions respectively. (F) ProSA-web, giving a Z-score of −5.67.

### Secondary and tertiary structure prediction of MEV and its refinement

The MEV protein intrinsically consist of diverse secondary structures. The protein was predicted to have 18.5% alpha helix, 29.08% beta sheet and 12.4% turns when analyzed by CFSSP:Chou & Fasman secondary structure prediction server (Figure 5 (C)). There were 23 amino acid residues constituting 6% of the protein is disordered structure. Additionally, Solvent accessibility estimation of amino acid residues provided 47% completely exposed followed by 24% medium exposed and 28% buried residues (Supplementary Figure 1). Structure validation of the refined 3D structure was performed using PROCHECK and ProSA-web (Figure 5 (D) & Figure 5 (E)). Assessment of the model with the Ramachandran plot verified 90.4% of residues in most favored region, followed by 9.3% on additionally allowed region and none on the outlier regions, which shows satisfactory overall quality of the model (Supplementary Figure 2). The quality of the construct was verified using ProSA-web which indicated the quality and prospective errors in the refined protein vaccine model with a Z-Score of −5.67 (Figure 5 (F)).

### Molecular Docking and simulations

The refined structure of MEV was docked with TLR4/MD2 complex (4G8A:AC) to examine adequate binding to immediate immune response. From the 20 predicted structures, top 10 solutions were refined further by using FireDock. The best-docked solution had binding energy of −37.63 kcal/mol (global energy), hydrogen bond (HB) contribution to global binding energy is −2.61 kcal/mol, and the contribution of the attractive and repulsive van der Waals forces to the global binding energy was −23.86 and 10.95 respectively (Figure 6 (D)) & (Table 7), which indicates a good binding affinity of vaccine with the TLR-4 receptor (Figure 6 (A), (B) & (C)). The RMSD plot of the MEV – TLR4 complex implies a sustained fluctuation until 0.5 ns after which the complex was in a steady condition at RMSD value of approximately 0.3 nm for the rest of the simulation (Figure 7 (A)). This shows good interactions between the MEV and TLR4 and entails stability of the complex. The RMSF plot of the MEV-TLR4 complex indicates the regions having higher flexibility in addition to its stability (Figure 7 (B)). Higher spikes in the RMSF plot indicates the high magnitude of flexibility of the residues which points at the modifications employed by the residues in respect to their interaction with TLR 4 and hence reinforces stability of the system with appropriate interactions. The plot of radius of gyration estimates the compactness of the MEV-TLR4 complex which is due to the molecular interactions between the MEV and TLR4 during dynamics (Figure 7 (C)).

**Figure 6:**
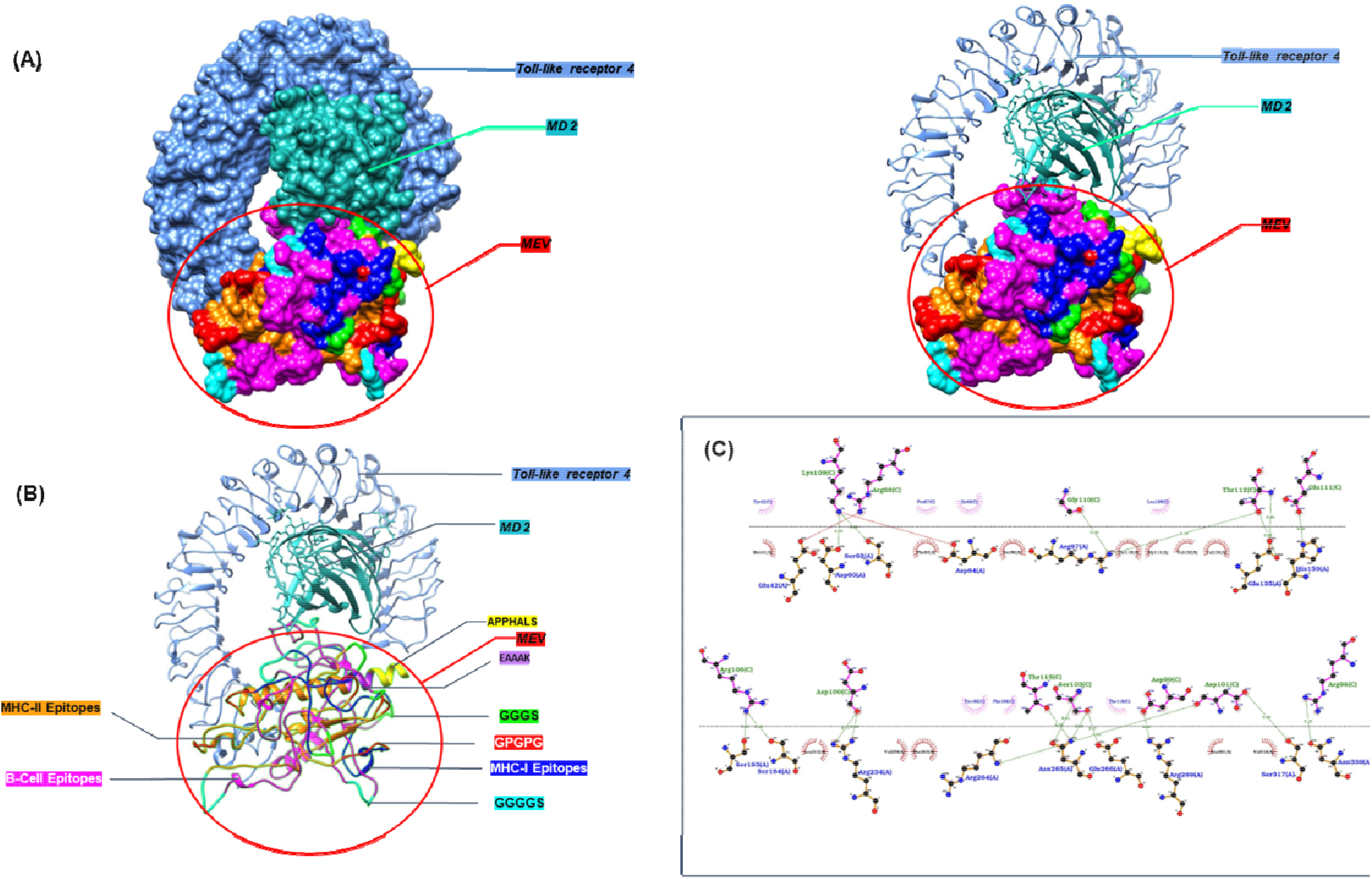
The interaction between the MEV and TLR-4 receptor as three- and twodimensional representations. (A) The docked complex of vaccine construct and its receptor TLR-4. (B) Various components of the vaccine construct, including intramolecular adjuvants (e.g., Leptospira surface adhesion (Lsa21), and APPHALS), linkers (e.g., EAAAK, GPGPG, GGGS, GGGGS), and B-cell and T-cell epitopes are shown as a docked complex with TLR-4. (C) Dimplot representation of hydrophobic and hydrogen bonds between the vaccine construct and TLR-4.

**Figure 7:**
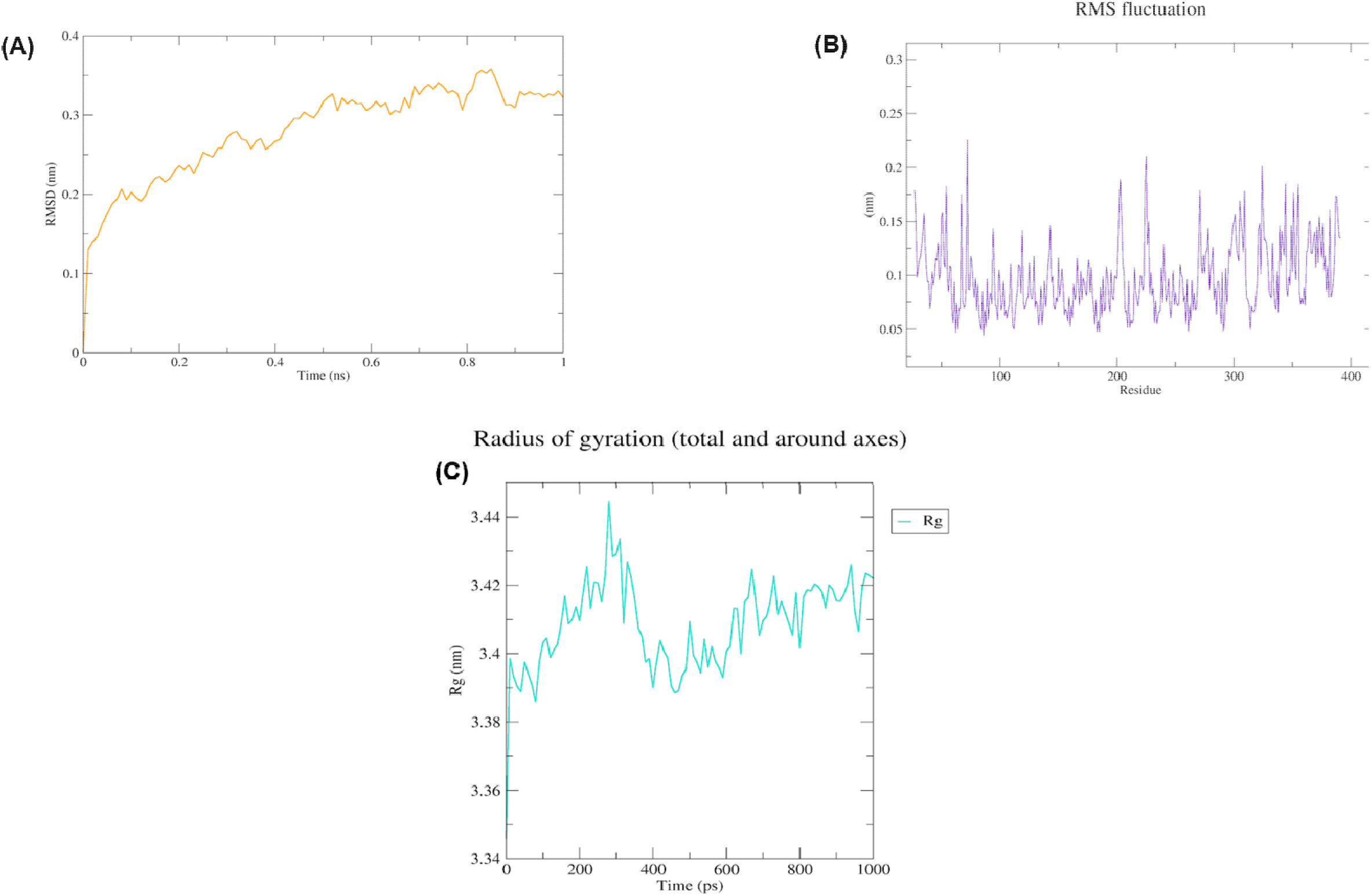
Molecular dynamics (MD) simulation of TLR4-MEV complex. **(A)** RMSD plot of the MEV-TLR4 complex emphasizing its dynamics and the stability, (B) RMSF plot of the MEV-TLR4 complex shows the flexibility of MEV-TLR4 complex, (C) Rg plot of MEV-TLR4 complex indicates the compactness of structure.

**Table 7:** Fire dock refined values of top 10 solution.

| **Rank** | **Solution**<br>**Number** | **Global**<br>**Energy** | **Attractive**<br>**VdW** | **Repulsive**<br>**VdW** | **ACE** | **HB** |
| --- | --- | --- | --- | --- | --- | --- |
| 1 | 6 | -37.63 | -23.86 | 10.95 | -1.29 | -2.61 |
| 2 | 3 | -30.79 | -16.57 | 6.23 | -4.94 | -0.84 |
| 3 | 9 | -8.32 | -18.67 | 5.42 | 16.40 | -3.53 |
| 4 | 4 | 3.76 | -31.86 | 8.90 | 18.13 | -4.79 |
| 5 | 7 | 12.50 | -1.25 | 0.00 | 0.69 | 0.00 |
| 6 | 5 | 166.78 | -33.11 | 260.80 | 3.75 | -6.00 |
| 7 | 1 | 233.68 | -23.68 | 324.58 | 11.26 | -5.69 |
| 8 | 10 | 438.86 | -39.73 | 647.01 | -2.80 | -3.23 |
| 9 | 8 | 1213.55 | -34.83 | 1588.11 | -2.58 | -3.43 |
| 10 | 2 | 3655.65 | -46.58 | 4717.55 | -18.99 | -4.70 |

### In Silico Immune Response Simulation

C-ImmSim immune simulator was used to produce a prompt in-silico immune response which was in accordance to the actual immune response produced. Both B and T -cell promiscus epitopes are present on MEV that can effectively trigger both humoral and cellular immune reactions, the production of IgG and IgM antibodies is triggered as a primary reaction (Figure 8 (A)), while the secondary reaction stimulates enhanced levels of IgG1 + IgG2, IgM, and IgG + IgM significant memory cell development (B-cell populations) antibodies (Figure 8 (B)). The IFN-γ concentration was high because of activation of IL-10 and IL-2, TH cell population are reported to be high during secondary and tertiary reactions which gradually decreases (Figure 8 (C and D)). Thus, these diverse immune reactions indicate that our vaccine construct is conceivable as a subunit vaccine and provide the basis for immunity against *Leptospira-associated* infections.

**Figure 8:**
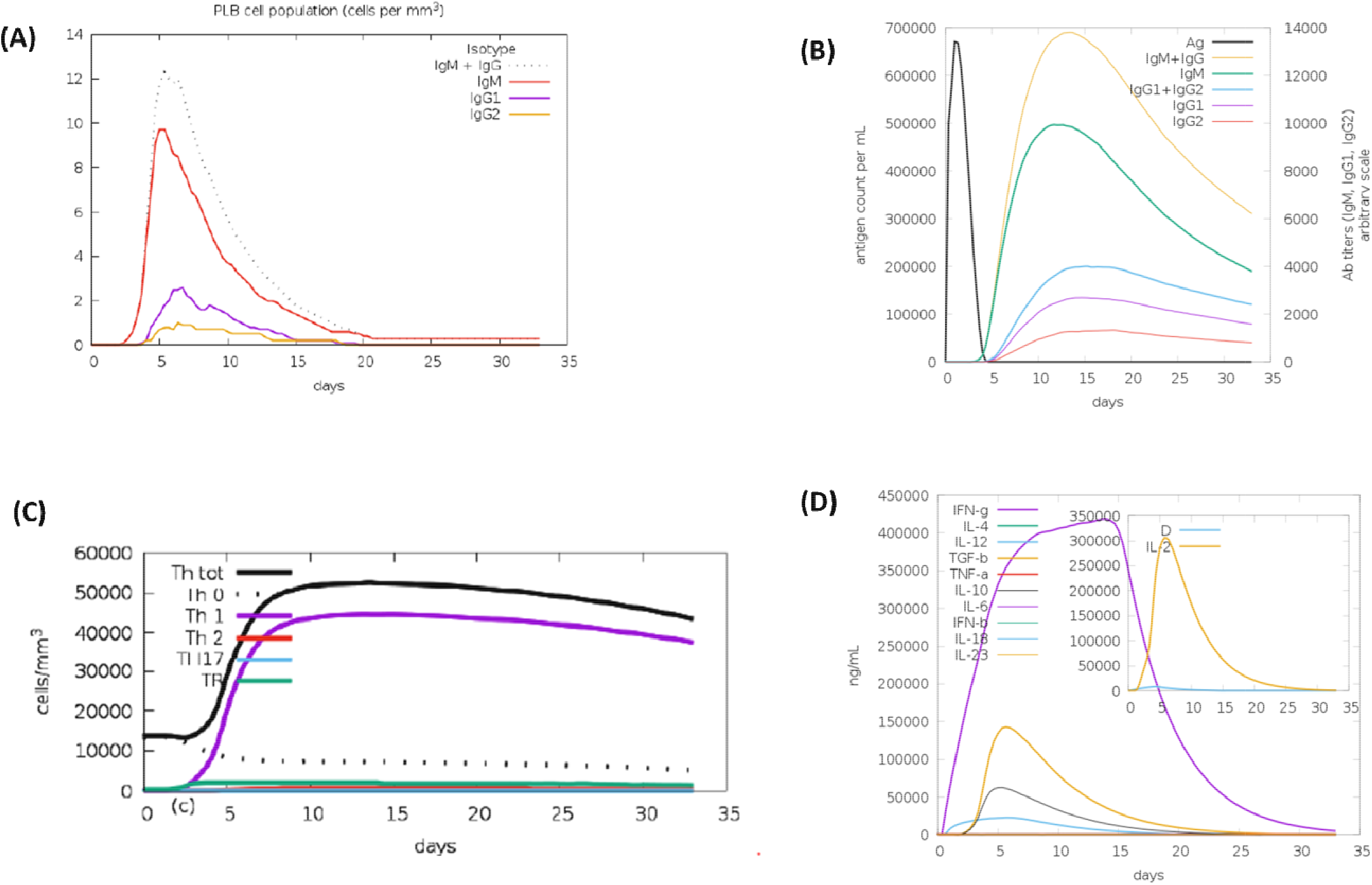
Immune simulation by C-ImmSim server. (A) Both IgG and IgM reached their maxima after MEV vaccine induction. (B) The immunoglobulins and the immunocomplex response to antigen (MEV) inoculations (black vertical lines); specific subclasses are indicated as colored peaks, (C) Evolution of Th cells with course of vaccination. (D) Induction of interleukins after vaccination with MEV.

### Optimization of codon and Insilco cloning

JCAT software was employed to reverse translate and optimize the codons of MEV to ensure it is highly expressed in the E. coli system. The optimized and reverse translated sequence was 1082 nucleotides. The CAI score calculated by JCAT is 0.98 for the given sequence, on the other hand GC content of the improved sequence was 33.05%; these scores stipulate that expression of the vaccine in E. coli K12 strain. BamHI and XhoI restriction sites were added to both ends of the vaccine gene. The vaccine genes were cloned into pET28a (+) plasmid using Snap gene software and the clone was 6.678 kb long (Figure 9).

**Figure 9:**
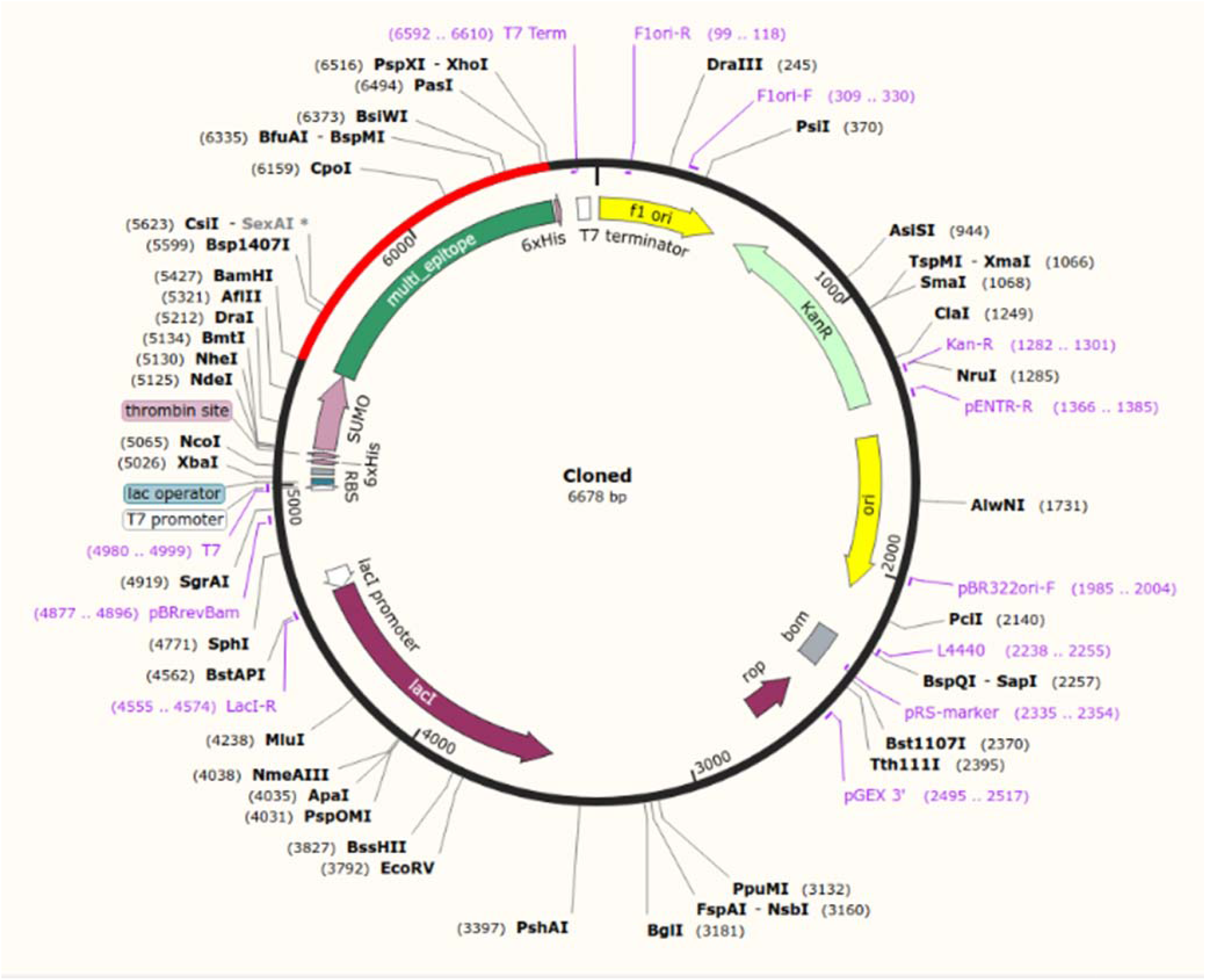
In silico fusion cloning of the MEV in pET28a (+) vector containing SUMO tag and 6X-His tag. The final vaccine candidate sequence was inserted into the pETite expression vector where the red part represents the gene coding for the predicted vaccine, and the black circle represents the vector backbone.

## Discussion

Prevention against the zoonotic disease *Leptospirosis* is still challenging even after availability of enormous immuno-informatics tools. Targeting soft-core or core genes of bacterial species could help in identifying better vaccine candidates to control the infectious disease (Aslam et al 2020). Conventional vaccine development approach is a laborious technique with low effectiveness and consumes a high cost in terms of money and time. In contrast, designing vaccine with the help of novel immuno-informatics approach is stable, safe, specific, more effective and less costly in terms of time and money (Rappuoli, 2010). Few prominent and widely used *Leptospirosis* vaccines in the market are multivalent inactivated whole-cell vaccine such as SPIROLET from France (Bashiru 2018), Vax-Spiral from Cuba (Martínez, R. 2004, Bashiru 2018), bivalent vaccine from china (Hu 2014) and Nobivac L4 from UK (BVA, 2016, Bashiru 2018). However, these vaccines (bacterins) has limitation in its application due to numerous associated drawback such as lack of cross-reactivity or protection to only closely related serovars (Grassmann, 2017), lack of long term immunological memory, requirement of regular boosters due to short duration of immunity and sometimes high reactogenicity (Sonrier 2000, Dellagostin 2011, Dellagostin 2017, Adler 2015, Adler 2011).

By utilizing the cutting edge novel immuno-informatics tools the soft core genome of pathogenic species of *Leptospira* was explored to identify essential core proteins, non-homologous, non-allergen to humans, were considered as vaccine targets. In our study the soft core genome of all sequenced pathogenic strains of *Leptospira* consisting of 2408 proteins were selected. Host homologs were excluded to avoid autoimmunity with hosts e.g humans (Chaudhuri 2014, He, Yongqun *et al*, 2010) which resulted 52 OMPs. All 52 proteins were screened for antigenicity by using Vaxijen v2.0 server (Doytchinova *et al* 2007) and 36 OMPs were predicted as highly antigenic in nature. Further we selected adhesin like proteins, as adhesins are virulence factors that allow bacteria to attach to host cells and are known to be good candidates for vaccine (Wizemann, *et al* 1999). Only 19 proteins which fulfill all the criteria of each step of pipeline (Figure 2) were selected as confident proteins with good adhesin properties. Subsequently these 19 OMPs were submitted for the selection of non-allergen proteins with host (Cui, Juan, et al. 2007). Only one Outer Membrane protein (WP_011671323) that shows significant match with known allergen “Sar s 1 allergen Yv5032C08” were excluded from the analysis (https://www.uniprot.org/uniprot/Q3L7L6). In addition to the above mention properties these proteins were screened for B and T cell epitopes as B and T Lymphocytes are responsible for generating the immune response after recognition of peptides presented on MHCI and MHCII by Antigen presenting cells (APCs) like macrophages and dendritic cells (Mazar Khan et al 2019). These epitopes have been previously identified by documented online tools from various vaccine targets of *Leptospira* including proteins like Lp11, Lp21, Lp22, Lp25, Lsa30,OMPL1, LipL 47, Lp35,Putative lipoprotein, some OMPs and Ton B proteins etc. The *Leptospira* soft core genome revealed that the proposed vaccine candidates covers a comprehensive number of high affinities of MHC-I and MHC-II alleles along with linear B-cell epitopes. Hence our proposed vaccine candidate consists a total of 69 epitopes were identified as promiscuous epitopes of which 29 epitopes binds to MHC-I reference allele and 40 epitopes binds to MHC-II reference alleles. Having a high number of epitopes predicted to bind a wide range of human HLA alleles does not necessarily correlate to high immunogenicity. All protein antigens cover more than 90 % population of World as well as South Asian countries. Studies have shown that the recognition of the peptide-MHC-I (pMHC) complexes have a more substantial impact on the immune response induction than the epitope diversity (Ishizuka et al.,2009; Kotturi et al., 2008). Finally, 6 Potential Vaccine Candidates (PVCs) were selected considering Immunogenicity, Antigenicity, Allergenicity, Protegenicity and Population coverage (Table 6). These PVCs are poise to induce both B-cell and T-cells responses. Reverse vaccinology approaches have been applied in several infectious diseases such as malaria (Tuju et al 2017), multiple sclerosis (Islam et.al 2020) and Plasmodium falciparum (Pritam et al 2019). Even though there have been several endevours to develop vaccines pertaining to specific diseases, the attempts were unfruitful.

For constructing MEV we have selected the best epitopes with good immunological properties in a way that it represented at least one epitope for MHC-I, MHC-II and B cell from each vaccine candidates. so that even if one gets cut other epitopes remain. The epitopes preferred on the basis of their hydropathy index in which epitope “IQSSILFLFVIFLVLFQTNLE overlapped with B-cell and TH2 (CD4+) epitopes with a hydropathy index of 2.08. All the epitopes were stitched with three amino linkers that are “GGGGS”, “GGGS”, “GPGPG” for B cell, MHC-I and MHC-II epitopes respectively. Linkers are introduced as an essential element in the MEV engineering to enhance folding, stabilization, and expression and certain studies suggest that GGGS linkers are superior to AAY for epitope-based vaccines (Yang, Y et al. 2015). The linkers used were G-rich which elevates the immunogenicity of the multi-subunit vaccine constructs (Yang Y. et al. 2015). The MEV solely cannot enhance superior immune response. Hence, it is coupled with an adjuvant which helps protect against infections, increases stability and durability of the vaccine formulation meanwhile extending the ability of the vaccine to induce a strong immune response without an allergic reaction towards human. Therefore, an adjuvant, “APPHALS” which was 7 amino acids long was combined with EAAAK linker at the N-terminal of the construct. The structural (i.e. secondary and tertiary) evaluation of the construct discloses spatial arrangement of vital protein constituents and provides exceptional assistance for the study of ligand interactions, protein functions and dynamics. The molecular weight of the MEV was observed to be ~37kDa which comes in the ideal range for a vaccine construct i.e. 30-60kDa. Further analysis confirmed the stability of the construct with an instability index score of 31.14 which is smaller than the stability threshold of 40.0. The MEV was recognised as antigenic with an antigenicity score of 1.308. The theoretical PI of the construct was calculated to be 4.92 which depicted the optimum pH at which the proteins exhibited no net charge (Shi et al, 2005). The sum of all hydropathy values were deliberated to a GRAVY value which revealed that the MEV was hydrophilic in nature. The assembly of the vaccine was found to be thermostable and also soluble upon overexpression. The integration of aforesaid properties conceives that the construct is biologically considerable. The reliability of the vaccine model was examined on the Ramachandran plot wherein majority of the residues fell in the most favourable region. The quality and perceptive errors of the construct was validated using ProSA-web server with a Z-score of 5.62 which correlates with the fraction of residues in favoured regions of the Ramachandran plot. The molecular docking studies bring to light that there were stable interactions between MEV and TLR4-MD2 complex and less energy (−37.63) was needed for proficient binding. The binding of MEV with Toll-Like Receptors 4 (TLR4) implies that it might play an important role in the innate immune response (Lorenz et al 2002). Molecular dynamics simulation of the MEV with the TLR 4-MD 2 complex displayed complete stability during simulation. Presence of B cell and T cell epitopes in the MEV theoretically should trigger humoral and cellular immune response. The vaccine exhibited a primary response with a substantial increase in the production of IFN-**γ**, with significant activity of IL-10 and IL-2. There was a magnified macrophage activity observed along with durable Th1-mediated immune response which is crucial in immune simulation. In addition, succeeding the primary reaction there was an increase observed in the active helper T-cell population. Active antibodies such as immunoglobulins IgM, IgG and their isotypes show an elevation in their number upon MEV immunization which could be included in isotype switching. Codon optimization is significant for higher expression since foreign gene expression may fluctuate in the host cell genome due to mRNA codon irregularity. The GC content and Codon Adaptation Index (CAI) value of our vaccine construct were 33.06% and 0.98 respectively which clearly depicts higher levels of expression within the E. coli K 12 strain. Restriction cloning of the pETite containing SUMO (Small Ubiquitin-like Modifier) tag and 6x-His tag i.e. SUMO pET28a (+) vector was executed to obtain possible vaccine gene and benefits both the solubilization and affinity purification of the recombinant protein because of its affluent polar and charged residues of vaccine construct (Biswal et al., 2015).

In summary, the propound construct could trigger an innate and adaptive immune response with promiscuous CTL, HTL and B-cell epitopes along with the TLR4 adjuvant. The results from the above mentioned analysis exhibited prospective scope for the development of a new Multi Epitope Vaccine for inducing optimum protective immune response against *Leptospira* infection. Nevertheless, advanced in vitro and in vivo investigations might be vital to reinforce the findings of this study.

## Conclusion

In comprehension, a novel **M**ulti **E**pitope **V**accine against Leptospirosis was designed using integrated vaccinomics approach which incorporates promiscuous antigenic epitopes capable of producing both cellular and humoral immune responses. In construction of the MEV, diverse functional peptides were stitched together with an adjuvant and linker sequences which aided in enhancing its potent immune activity. Investigative analysis of the physicochemical properties and structural corroboration helped increase the efficiency of the vaccine construct. The construct exhibited desired characteristics and established interactions with the TLR 4-MD 2 complex which were validated through means of docking and simulation studies. Codon based cDNA of the MEV is anticipated to show high expression in the mammalian host cell line. Since the MEV is designed from antigens identified by comparative genomics and reverse vaccinology of various pathogenic serovars it is likely that it can induce cross protective immunity. However, experiments involving evaluation of immune response and protective efficacy against infection in relevant animal model is required to prove this prediction.

## Acknowledgements

This work was supported from DBT grant (BT/PR21430/ADV/90/246/2016) funded to SMF from Department of Biotechnology, Ministry of Science and Technology, Government of India. Financial support from NIAB core funds to SMF and SA is duly acknowledged. The authors would like to thank Director, NIAB for providing all the facilities and his constant support and encouragement.

## Author’s contributions

SA and SMF conceived the idea and designed the study. MA, MK and SA performed the analyses. RS provided critical insights and suggestions in various parts of the data analysis. MA and MK wrote the initial draft and SMF and SA editing the manuscript. All authors read and approved the final version of manuscript.

